# Frank-Starling law and Bowditch phenomenon may have the common theoretical grounds

**DOI:** 10.1101/069609

**Authors:** Yuri Kamnev

## Abstract

The relationship between two different linear dimensions of the chamber (ventricle) and the two respective values of the force of contraction (which are applied to the imaginary piston in order to accelerate the initial venous inflow) is deduced as the ratio of forces which is equal to the ratio of squares of linear dimensions of the chamber; the equation is valid when the durations of both contractions (systoles) are identical. The relationship corresponds to Frank-Starling law (the right limb of parabola can be approximated to the direct proportionality of the law). When the durations of systoles are different the ratio of forces is equal to the inverse ratio of durations of systoles; the inverse proportionality permits to interpret the Bowditch phenomenon by means of the ascending asymptote of hyperbole. Stepwise shortening of systole is impossible due to extremely narrow range of variable (duration of systole), hence the shift of variable can be realized only as a leap; this leap is accompanied by the enormous rise of function (force of contraction) which can be accommodated to several contractions. Homoiometric regulation can be considered a safety device (presumably in the form of paroxysmal tachycardia with the shortened systole) in the case when heterometric regulation lacks to produce the adequate force of contraction in response to excessively distended chamber.

---

Heterometric regulation (Frank-Starling law) and homoiometric regulation (Bowditch phenomenon) belong to the same phylogenetic level that justifies the attempt to formulate both of them in some general form and look for theoretical model of combination.

## Methods

Imagine the pipe of the constant diameter and the liquid moving in it with volume flow rate Q. The flow denotes the venous flow that lacks enough energy (low pressure flow) to overcome the resistance ahead, and it is the role of the pump to supply the flow with some extra-energy by means of the force F. Thus, the basic speed is Q (or linear velocity U if we deal with the travel of some solid mass) and this volume flow rate Q must be accelerated by means of application of the force F to the piston that mystically appears inside the pipe. Then we must mark out the active portion of the pipe: at the beginning of it a piston suddenly appears, it pushes the flow forward up to this or that stop-point; the length of this or that distance the piston covers (*l*_1_ or *l*_2_) corresponds to this or that end-diastolic volume of the ventricle.

Instantly we come across the following problem. The piston moves the columns of liquid *l*_1_ or *l*_2_ (that corresponds to the displacement of masses or *m*_1_ or *m*_2_) but these columns of liquid are connected to the continuation of the column of liquid that fills the pipe ahead (so to say, to the column of blood in the artery). Hence, the piston applies the force F to some huge masses but not to ffl] or *m*_2_. Nevertheless, the analogous process takes place when the semilunar valve is opening (isometric phase of contraction changes to isotonic) and maybe it is pertinent to represent masses as *m*_1_ + const and *m*_2_ + const. However, such an addition is not accurate if the diameter of the continuation of the pipe is variable. Consequently, our model matches to the case when the artery is short (factually of the constant volume) and it ejects blood into lacunae; the assumption is relevant because heterometric and homoiometric regulation is evolutionally most early type of controlling of the circulation. Therefore, we may neglect the constant volume inside the artery or may consider its invariable presence.

By the way, the virtual inner piston can be realized by the roller of peristaltic pump, - but this roller must be able to cover different distances (in contrast to roller of the ordinary peristaltic pipe).

## Results

Thus, we are going to compare two cardiac cycles of the same duration (i.e. rhythm is rigid); duration of systole *t_syst_* in both cases is identical, i.e. it takes the equal time intervals for the piston to travel along *l*_1_ or *l*_2_. Duration of diastole in both cases is also identical (as far as rhythm is rigid and duration of systole is constant) but the volume flow rates *Q*_1_ and *Q*_2_ are different, - that is why *l*_1_ and *l*_2_ are different (as if the ventricle is more distended in case with *l*_2_, for example).

Now let us scrutinize the ejection. According to Newton second law:

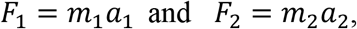

where *F*_1_ and *F*_2_ denote the forces which are applied to the piston during the systole in order to move *m*_1_ and *m*_2_, respectively (i.e. to move the masses of blood containing in the distended ventricles).

According to 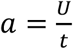 we may write:

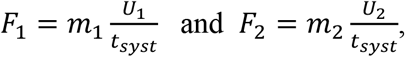

and then equate both formulae because *t_syst_* is standard:

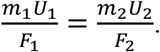

Now let us replace the masses by densities and volumes due to *m* = ρ*V*; after substitution we achieve the equation:

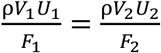

which can be transformed to proportion:

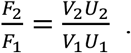

The final transition to hydrodynamic terminology can be done by representation of *VU* as *Ql* according to the equivalence 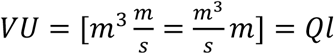:

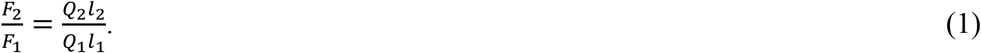

Our model where the diameter of the pipe is standard permits us to find the value of *Q*_2_ by means of the relation of piston travels 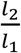 applied to the value of *Q*_1_:

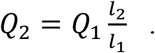

After substitution of *Q*_2_ into the proportion (1) we can get rid of the volume flow rate in 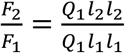 and finally conclude:

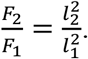

Numerically, for example, when the length of imaginary fiber was elongated from 5 to 7 the two-fold increasing of F appears as the response to such elongation:

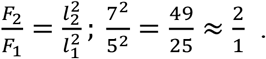

Miscellaneous diagrams picturing the dependence of tension (or the contractile force) and the length of myocardial fiber, - cited as data of different experiments, - demonstrate non-linearity of the linkage. If roughly: the shallow slope at small lengths and steeper slope at larger lengths, i.e. the operating range of Frank-Starling curve resembles very much the right limb of parabola (Fig.1). The curvature can be hardly discerned (referring to experimental data) but the steepness of such direct proportionality (that claims significant numerical coefficient before the argument) can be approximated by the respective region of the appropriate parabola.

**Fig.1.**
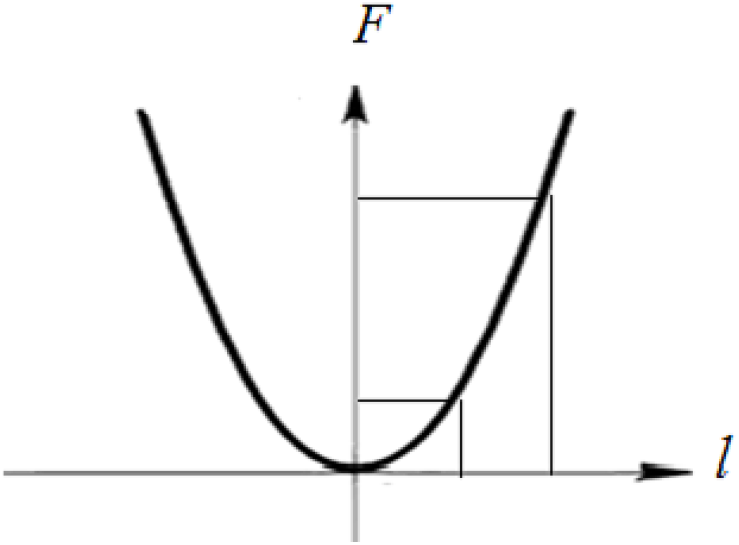
The right limb of parabola as the description of Frank-Starling law: the region of variable (distension of the muscle or linear variation of the chamber) and the region of function (force of contraction) are shown.

Now let us repeat the above deduction while modifying the conditions. What is in common: the duration of diastoles are equal. What is different: firstly, the volume flow rate Q is the same in both cases and, consequently, the end-diastolic volumes are identical (*l*_1_ = *l*_2_, if in terms of linear dimension of the pipe), as well as masses of liquid, *m*_1_ = *m*_2_ which fill the equal portions of the pipe; secondly, the durations of systoles are different *t*_*syst*1_ ≠ *t*_*syst*2_, and that results in the difference of the heart rate.

Let us begin the deduction in mechanical terms (as it was done before): we shall use masses, *m*_1_ and *m*_2_, like solids pushed by the pistons and linear velocities of pistons denoted as *U*_1_ and *U*_2_.

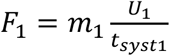 and 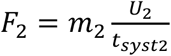, but *m*_1_ = *m*_2_ and, consequently, 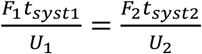.

The equation can be easily transformed to proportion:
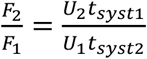, where *U*_1_ = *U*_2_, that corresponds to the condition of the standard value of Q.

Hence, 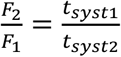.

The contractile force as the function of the duration of systole demonstrates the inversely proportional bond. The inverse proportionality is a non-linear function with very contrast extremes (hyperbole) and such function can be applied to physiological process only as a restricted operating range which is chosen according to observations. Physiological variation of the duration of systole (assumed to be the range of variable) is quite narrow and physiological variation of the contractile force (assumed to be the range of function) is very broad, - as it follows from measuring. The above variations conform to the region of variable and to the region of function of the section of hyperbole shown on Fig.2. Let us compare the restricted parts of curvatures on Fig.1 and Fig.2. The approximation of the right limb of parabola as direct proportionality, - or taking this section of square function as it is, - supposes some stepwise increasing (decreasing) of function as a response to the stepwise increasing (decreasing) of argument; the approximation of the ascending asymptote of hyperbole is something like a leap: stepwise increasing (decreasing) of argument is almost impossible (undetectable) if you want to achieve the stepwise decreasing (increasing) of the function. The leap-like shift of variable (duration of systole), - in order to reach some smaller value, - results in the enormous growth of force of contraction. The quick growth of the force is accommodated to several consecutive contractions, - which demonstrate that the slope of the function exists nevertheless.

**Fig.2.**
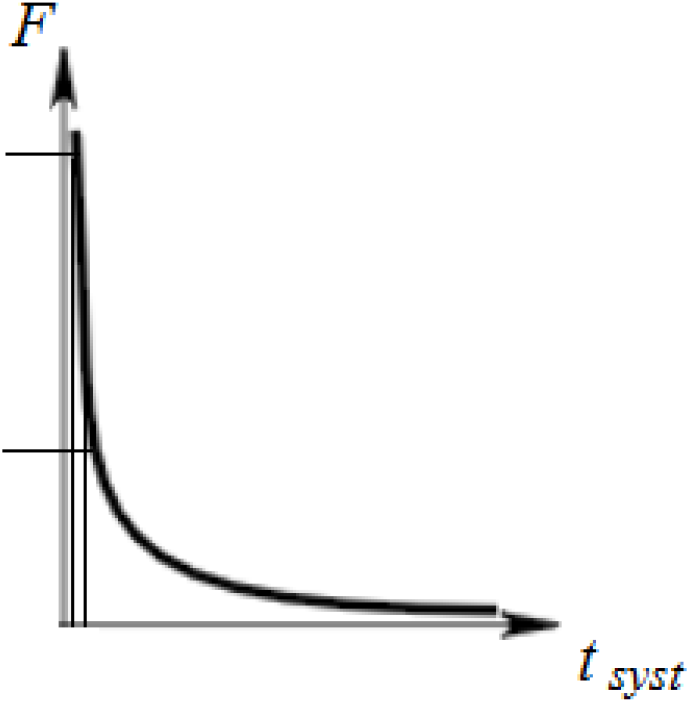
The ascending asymptote of hyperbole as the description of Bowditch phenomenon: the region of variable (duration of systole) and the region of function (force of contraction) are shown.

The shortening of systole means the shortening of the cardiac cycle (ventricular cycle, if strictly) and the quick growth of contractile force as a response to both shortenings can be realized as the specific adaptive reaction of the new type – which can be called paroxysmal tachycardia. Let us determine (presume) the situation when the adaptive reaction of such type is indispensable. Imagine the early period of evolution of circulation when the pace maker (as an inducer of different rhythms) is not exist yet and the control of circulation is carried out by Frank-Starling law on the basis of the rigid rhythm. At some extraordinary situation the volume flow rate Q becomes very high and causes the distending of the ventricle that exceeds maximal value of *l*_2_; in other words, there is no the respective value of *F*_2_ at Frank-Starling function. The way out is to create such value of *F*_2_, - or, more probably, to create the value that exceeds significantly the expecting one. It is time to switch on mechanism of urgent unloading, i.e. to shorten the duration of systole in order to achieve abrupt increase of contractile force. The shortening of systole influences slight acceleration of pulse due to the shortening of cardiac (ventricular) cycle, and this can trigger (for the system with rigid rhythm) the generation of another, more rapid, rhythm, i.e. the paroxysmal rhythm. Let us turn again to the operating range of the ascending limb of inversely proportional function (Fig.2). The shift of argument (on abscissa) closer to zero can be accomplished only as a leap; the range of function, being displayed to the long distance of ordinate, is able to reveal the stepwise increasing of the value if the carrier of the value has the peculiarity of being divided: the contractions are separated one from another, so the increasing value of contractile force may appear gradually at several contractions and then reach some steady maximum.

(It is interesting that the mechanism of unloading of excessively distended ventricle, i.e. paroxysmal tachycardia at the evolutionary early type of circulation, possesses the intrinsic feedback that switches-off this rapid rhythm. The abrupt acceleration of pulse can not be affected by shortening of the systole only; inevitably the duration of diastole will be involved, - and it will be shortened either. If so – the filling of the ventricle will be diminished in spite of the constantly high Q. The value of distention of the ventricle will return to the range of variable and the value of the newly adequate *l*_2_ will correspond to the respective *F*_2_ according to Frank-Starling law. The reappearance of active Frank-Starling regulation may signal to disconnect the device that has triggered paroxysmal rhythm and shortens the systole.)

The Bowditch phenomenon [1] looks very much like the triggering of the above mechanism by means of changing the initial frequency of stimulation to more frequent. Stimuli that are more frequent play the role of paroxysmal tachycardia which causes the shortening of systole and, as a feedback, the muscle switches on the realization of function at the ascending limb of hyperbole. Therefore, the staircase phenomenon originates from the function (inversely proportional) that was deduced together with another function – square function, the section of which can be approximated as almost direct proportionality of Frank-Starling law.

The combination of Frank-Starling regulation with homoiometric reaction is the combined mechanism of the real regulation with the safety device preventing from the excess of diastolic distending due to the inflow exceedingly high. Such system is relevant when heterometric regulation is supported by the rigid rhythm only and no controlling reflexes exist yet. The next step of the evolution of circulation (when the rhythm becomes changeable and the basic reflexes appear together with pressure detectors) has solved the problem of overloading of the ventricle by means of another approach to the problem. More advanced system tries to avoid the extreme by means of early switching on the regime that was substituted instead of the homoiometric reaction. The mathematical model of circulation controlling - which relates to the level of evolution mentioned above – is available [2].

